# Genetic Novelty and Short-Term Evolutionary Dynamics in *Drosophila yakuba mayottensis*

**DOI:** 10.1101/2025.06.10.658671

**Authors:** Brandon A. Turner, James E Titus-McQuillan, Amir Yassin, Rebekah L. Rogers

## Abstract

Understanding how genetic novelty arises and contributes to population divergence is central to evolutionary biology. On short evolutionary timescales, the relative roles of new mutations and standing variation remain underexplored among structural variation. Here, we compare the contributions of chromosomal rearrangements, tandem duplications, and single-nucleotide polymorphisms (SNPs) to early genomic differentiation in a recently colonized island population, *Drosophila yakuba mayottensis*, from the island of Mayotte. By comparing the relative impacts of these mutation classes, along with the involvement of transposable elements (TEs), we assess whether certain mutations are more likely to contribute to rapid divergence. We show that TEs are disproportionately involved in the formation of novel structural variants shortly after colonization, particularly through direct insertions. Our results suggest that TE-associated rearrangements may bypass the clocklike pace of SNP accumulation and divergence, where the contributions of coding SNPs, tandem duplications, and TE-facilitated ectopic recombination are constrained to standing variation on short timescales. Although TEs can drive the rapid emergence of novel variants, their full contribution to adaptation likely depends on evolutionary context and time since colonization. Together, our findings reveal time-dependent mutational dynamics and highlight the importance of structural variation, particularly TE-associated rearrangements, in shaping early genomic divergence following habitat shifts.

## 2 Introduction

The relative importance of standing genetic variation versus new mutations plays a central role in shaping the tempo of genomic change. Standing variation provides a pool of preexisting alleles that can respond immediately to new selective pressures or environmental shifts, often enabling rapid adaptive responses [1–4]. In contrast, new mutations must first arise and subsequently overcome stochastic loss, a process that typically introduces a delay in generating novel variants [1, 5]. For populations experiencing sudden habitat shifts or environmental stress, this distinction becomes critical. Evolutionary responses from standing variation may occur almost instantaneously, while new mutations might only contribute over longer periods unless mutational bursts accelerate their availability [6, 7]. Understanding which sources of genetic novelty contribute to variation on short evolutionary timescales can help reveal the mechanisms that allow populations to generate novel mutations quickly under shifting selective regimes [1, 8].

The timeline to generate new mutations is expected to be long for SNPs, even in species with large population sizes [9, 10]. However, other types of mutations, like new TE-induced mutations, may violate the assumptions of the molecular clock [11, 12]. If these alternative modes of genetic change modify genes more rapidly than SNPs, they may provide mechanisms of variation that are more relevant on short evolutionary timescales [12,13]. If new mutations from these mechanisms arise as slowly as single base pair changes, then standing variation remains the primary reservoir of change as populations invade new environments [2, 4, 14]. However, if TE-induced rearrangements and structural variants arise more quickly than SNPs, then such mutations may play a critical role in adaptation by allowing populations to bypass long mutational waiting times that typically constrain evolutionary responses [14,15].

### 2.1 Structural Variation and SNPs in Mutational Dynamics

Structural variation, including chromosomal rearrangements and tandem duplications, is increasingly recognized as a major contributor to mutational dynamics (that is, the timing, rate, and spectrum of mutations) particularly in populations experiencing rapid environmental or demographic shifts [16–18]. While SNPs act on individual base pairs, structural variation involves large-scale mutations that reshape genomic regions, influencing multiple genes or regulatory elements simultaneously [1, 19]. These structural changes can accelerate the introduction of genetic novelty by altering genome architecture, reshuffling regulatory networks, and creating novel gene combinations [20–23].

Both SVs and SNPs contribute to genetic differentiation, but they operate on different temporal and mechanistic scales [1, 24, 25]. Nonsynonymous SNPs that change amino acid sequences are among the most well-studied mutational changes and have been documented to contribute to population differentiation and adaptive evolutionary change [1, 26, 27]. However, SNPs typically accumulate incrementally, contrasting with the potentially rapid genomic restructuring observed in SVs, which can affect multiple genes or regulatory regions simultaneously [9, 28].

One key mechanism underlying this structural change involves the activity of transposable elements (TEs). Recent studies have highlighted the role of TEs in mediating structural variation, acting as catalysts for chromosomal rearrangements through mechanisms such as ectopic recombination or the direct insertion of duplicated sequence adjacent to the element [29, 30]. TE-mediated structural variants can arise in bursts, introducing new genomic architecture that circumvents the slower mutational dynamics of SNP accumulation [16, 31, 32]. While many TE-induced mutations are neutral or deleterious, a subset may persist and contribute to divergence by reshaping genome architecture and altering regulatory networks [33–35].

### 2.2 Mutational Constraints and Adaptive Trajectories

While adaptation requires beneficial mutations, the emergence and persistence of new mutations are often constrained. They must first occur, escape stochastic loss, and subsequently reach appreciable frequencies within a population [1, 2, 5]. These hurdles, especially in the early stages of adaptation, mean that populations often cannot access the full range of potentially advantageous changes that theoretically exist, given time. This is particularly relevant on short evolutionary timescales, where rare beneficial mutations may not arise quickly enough to contribute meaningfully [1,6]. Under such conditions, large-effect mutations, those that could shift adaptive trajectories most dramatically, may remain inaccessible, either because they are slow to appear or because they are lost early due to stochastic processes. This includes effects described by Haldane’s sieve, which can limit the fixation of recessive beneficial mutations [36]. If mutation constrains evolutionary walks, the initial steps of adaptation will be limited to standing variation [2, 8].

If beneficial mutations arise quickly, however, populations can explore genotype space more freely during adaptation [14, 37]. Under Fisher’s model of adaptive walks [14, 38], mutations of large effect are expected to fix early, followed by smaller-effect mutations that fine-tune adaptation. But this model assumes all beneficial mutations are equally accessible. When mutation rates vary or some classes of mutations are constrained, populations may instead follow suboptimal paths shaped by the limited variants available in standing variation [14, 15]. Populations free from mutational constraint may ultimately reach higher fitness peaks than those limited to pre-existing genotypes. In such cases, convergence at the mutation level may be more common among shared standing variants, while new mutations are more likely to be rare, heterogeneous, and less likely to produce identical outcomes with mutation level convergence [37, 39].

Different mutation types, such as SNPs, tandem duplications, chromosomal rearrangements, and TE-mediated variants, arise at different rates and through distinct mechanisms. While SNPs often accumulate gradually over time [9, 10, 28], structural variants and TE-driven mutations may occur in irregular bursts [11, 31, 32]. These bursts can violate the assumptions of a uniform molecular clock, leading to uneven contributions of different mutation classes to adaptive trajectories. As a result, the tempo and direction of adaptation may depend not only on selection but also on the specific mutational mechanisms and their emergence times across the genome.

### 2.3 Ecology of *D. yakuba* on Mayotte

To address these open questions in evolutionary genetics, we have searched for systems that experienced recent shifts in selective pressure. The island of Mayotte provides a natural experiment for studying mutational dynamics on short evolutionary timescales. On Mayotte, the population of *D. yakuba* (*Drosophila yakuba mayottensis*) is relatively young, diverging from mainland populations between 10,000 and 28,000 years ago [40]. This short divergence time presents a unique opportunity to assess the role of new mutations versus standing variation in generating structural variants and SNPs. The population’s dietary specialization on toxic noni fruit (*Morinda citrifolia*) may have further exposed pre-existing structural variants or driven new mutational events, highlighting how ecological shifts can influence mutation rates and genomic architecture [41, 42].

Understanding the relative contributions of these mechanisms in newly colonized populations like Mayotte’s *D. yakuba* can provide insight into how mutation dynamics are shaped by selection, stochastic processes, and mutational constraints.

## 3 Results

### 3.1 Prevalence of SNPs, SVs, and TEs

To understand how different classes of mutations contribute to genetic differentiation following recent habitat shifts, we characterized the prevalence of chromosomal rearrangements, tandem duplications, and SNPs in the Mayotte *D. yakuba* population and compared them to the ancestral mainland *D. yakuba* population. By quantifying their genomic distributions and allele frequency patterns, we aimed to assess how structural variants (SVs) and SNPs differ in their evolutionary dynamics following island colonization.

Chromosomal rearrangements were identified by analyzing abnormally mapping read pairs from Illumina paired-end reads, revealing a total of 7,021 unique rearrangements. Among them, 1,179 rearrangements were detected in 18 strains from the Mayotte population, and 6,345 rearrangements in 19 strains from the ancestral mainland *D. yakuba* population. Furthermore, we identified a total of 701 tandem duplications across both populations, with 462 present in Mayotte and 492 in the ancestral population. Notably, the number of rearrangements identified in the mainland population, particularly singletons, is higher than in our previous study due to the use of an updated *D. yakuba* reference genome (*Prin Dyak Tai*18*E*2 2.1) [16]. This increase reflects improved detection sensitivity and resolution of structural variation. In addition, minor differences in reference quality and PCR duplicate levels across samples may influence the absolute counts of low-frequency rearrangements, especially singletons, though such effects are unlikely to drive broad patterns of population-level differentiation.

**Figure 1:**
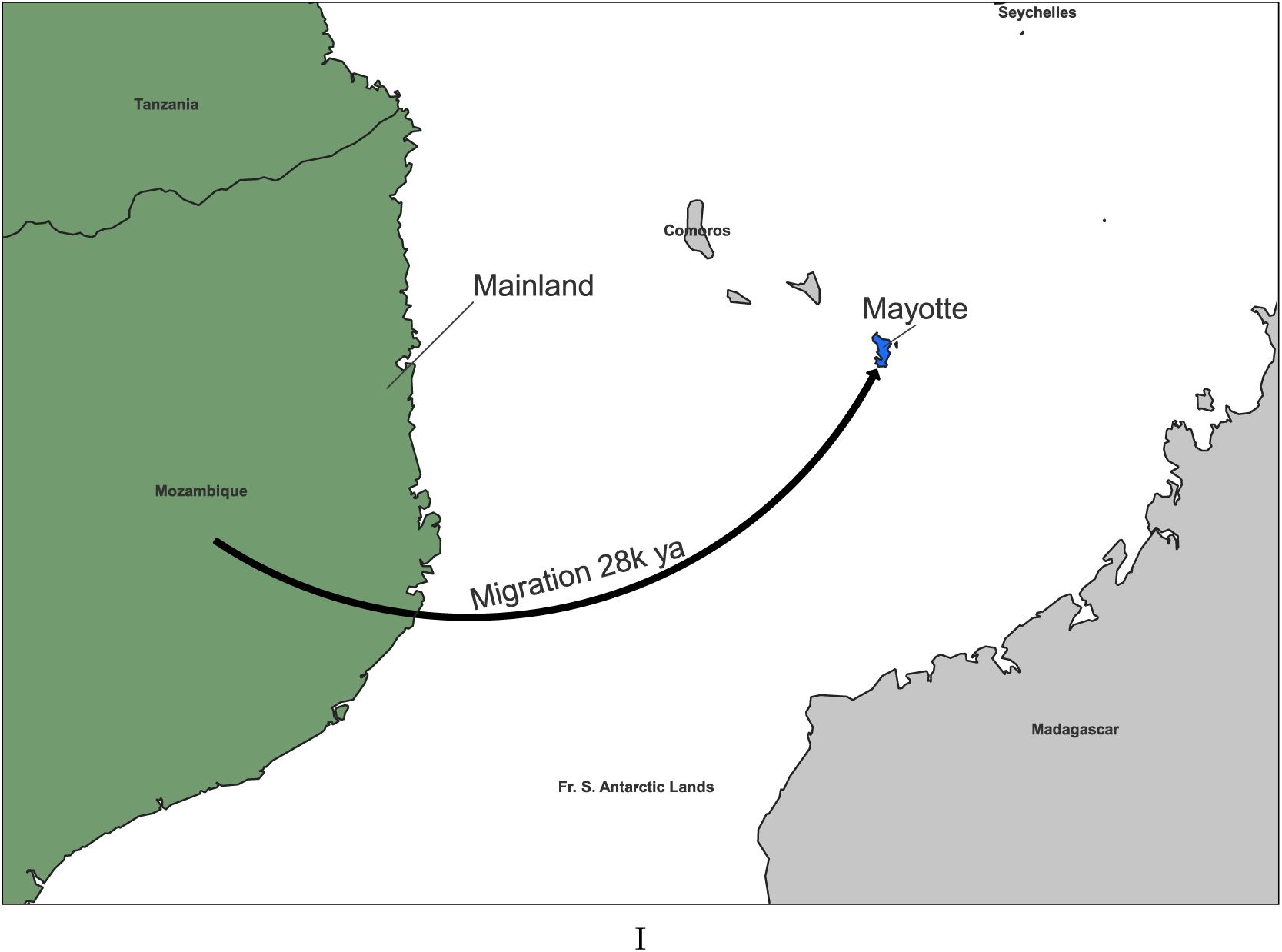
Map depicting *D. yakuba* habitat shifts from mainland Africa to São Tomè at two different time points.

Our analysis revealed a strong association between structural variants (SVs) and transposable elements (TEs). In Mayotte *D. yakuba*, 57.9% (5,132/8,850) of rearrangements were associated with TEs. Expanding this analysis to tandem duplications, 35.2% (2,880/8,178) of tandem duplications had at least one duplication breakpoint associated with a TE.

SNPs were identified using bcftools and SNPGenie (see Methods). We identified 3,310,289 SNPs genome-wide in the Mayotte population, with 247,423 SNPs falling within gene-coding regions. Using SNPGenie, we further identified 38,544 synonymous and 157,449 nonsynonymous SNPs within coding sequences. Having established the prevalence of these SNPs, we next evaluated their allele frequency distributions across populations to determine whether shifts in allele frequency reflect distinct mutational dynamics following island colonization.

To evaluate allele frequency distributions, we applied derived site frequency spectrum (SFS) analysis, correcting for uneven sample sizes between populations (see Methods). Among rearrangements, TE-associated variants were disproportionately represented among mutations at high allele frequencies and show that rearrangements significantly differed from expectations established by within-genome neutral SNPs (Kolmogorov-Smirnov, *D* = 0.721, *p <* 3.50 × 10*^−^*^14^). This trend, both in TE skewed allele frequency and deviations from neutral expectations, is also seen among tandem duplications (Kolmogorov-Smirnov, *D* = 0.739, *p <* 2.20 × 10*^−^*^16^).

SNPs in gene-encoding regions were also assessed for their allele frequency distributions. In the Mayotte and mainland populations, we identified 24,126 and 75,351 nonsynonymous mutations, as well as 42,694 and 95,551 synonymous mutations, respectively. These allele frequency distributions also showed significant differences from neutral expectations in both populations (Kolmogorov-Smirnov test, Mayotte: *D* = 0.3, *P* = 0.3356; Mainland: *D* = 0.65, *P* = 0.00027).

### 3.2 Rapid divergence between island and ancestral mainland *D. yakuba*

Building on the observed prevalence of SNPs, SVs, and their TE associations, we next evaluated how these variants contribute to shifts in allele frequencies between Mayotte and mainland populations. By quantifying these shifts, we aimed to determine whether mutation-driven dynamics influence genomic differentiation following island colonization. To quantify these shifts, we analyzed chromosomal rearrangements, tandem duplications, and SNPs within gene-coding regions using population branch statistics (PBS) and pairwise allele frequency differences (Δ*p*). PBS enabled the detection of loci with extreme allele frequency changes, indicative of underlying mutational dynamics. Pairwise Δ*p* comparisons further highlighted specific mutations contributing to this differentiation.

Using Δ*p*, we identified 74 of 8,850 chromosomal rearrangements (0.84%), 177 of 8,178 tandem duplications (2.16%), and 16,086 of 247,423 coding SNPs (6.50%) as significantly differentiated. Building on this, we utilized PBS to pinpoint specific loci driving these frequency shifts.

PBS analysis across Mayotte and mainland populations identified regions with pronounced allele frequency changes, many overlapping with known inversion breakpoints [43] in the Mayotte population. These peaks likely reflect mutation-driven structural changes facilitated by TE activity or regional mutational hotspots [41]. PBS analysis across Mayotte and mainland populations identified regions with pronounced allele frequency shifts, many overlapping with known inversion breakpoints in the Mayotte population [43]. These peaks likely reflect mutation-driven structural changes facilitated by TE activity or regional mutational hotspots. One notable region, an X-linked inversion spanning 8.68Mb-9.23Mb, exhibited concentrated structural and sequence-level differentiation [41,43]. Within this window, RepeatMasker [44] analysis identified 372 insertions from the SAR2-DM transposable element family clustered in a 55-kb interval (8.85Mb-8.91Mb), alongside 58 significantly differentiated coding SNPs. The co-occurrence of TE insertions and differentiated SNPs within this region suggests mutational hotspots driven by TE-mediated instability. While the exact mechanisms remain unresolved, these findings are consistent with TEs accelerating structural variation and allele frequency shifts in isolated populations.

**Figure 2:**
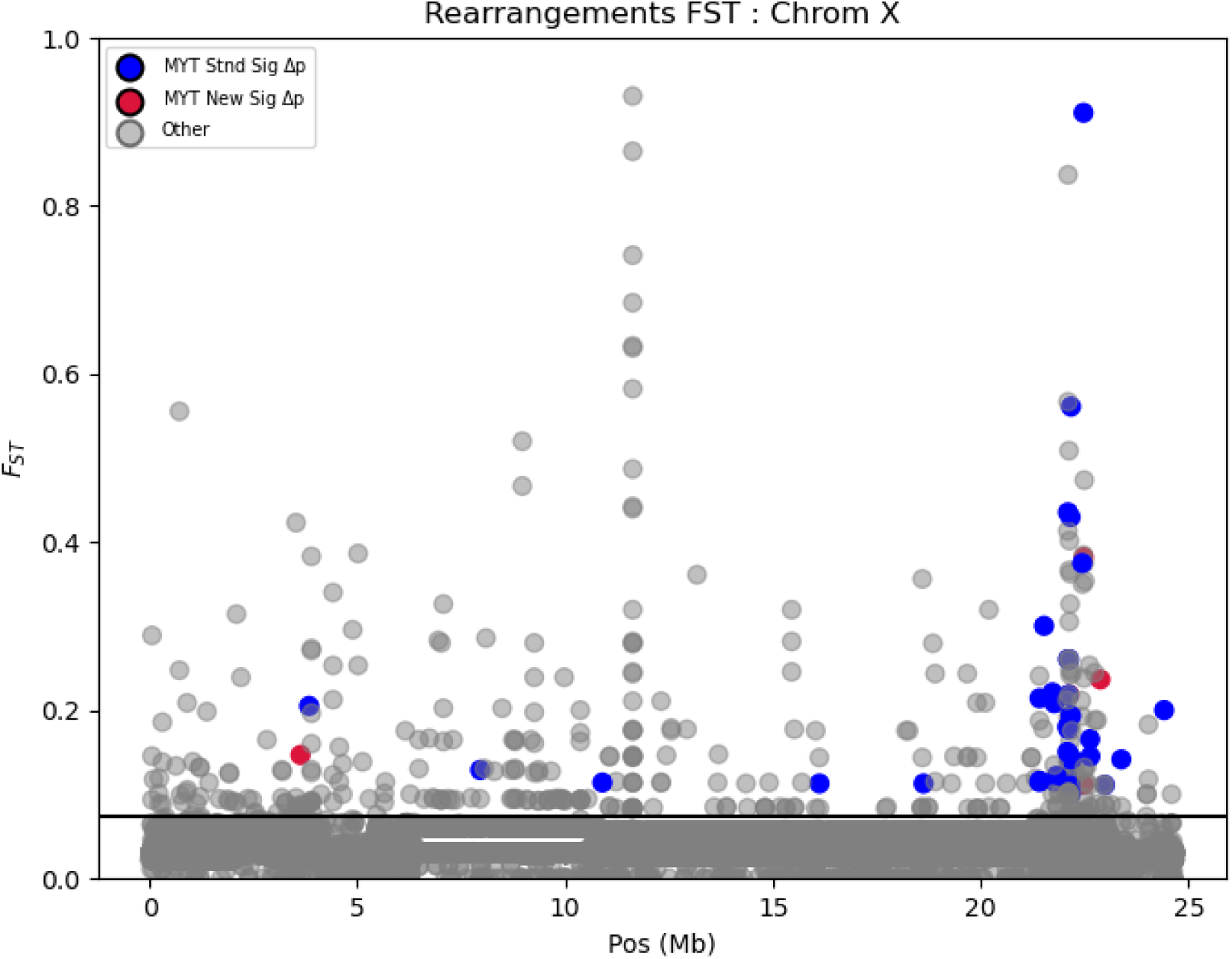
Multiple differentiation metrics (FST and Δ*p*) for chromosomal rearrangements on the X chromosome. Plotting FST values for all rearrangements, and then highlighting those with significant allele frequency differences (Δ*p*) for rearrangements disproportionately observed in the Mayotte *D. yakuba* population. Blue points represent significantly differentiated standing variation on Mayotte, and red points represent significantly differentiated new mutations on Mayotte. The horizontal black bar establishes the Δ*p* significance threshold. A concentration of significant standing variants is visible, consistent with the enrichment of ectopic recombination events among differentiated rearrangements.

Beyond localized inversion regions, structural variants were broadly distributed across the genome, with chromosomal rearrangements and tandem duplications appearing in both structurally complex and collinear (structurally conserved) regions. This widespread presence suggests that mutational activity is not limited to recombination-suppressed regions, reflecting a broader landscape of genomic instability. Transposable element insertions were similarly distributed, reinforcing the idea of continuous mutational activity across varying recombination environments. These patterns highlight the pervasive influence of TEs in promoting structural variation, potentially accelerating the emergence of novel variants during early island colonization.

### 3.3 Feature Association from Different Mutational Classes

To investigate the genetic mechanisms underlying differentiation, we analyzed the association of chromosomal rearrangements, tandem duplications, and coding SNPs with TEs. Given the role of TEs in mediating ectopic recombination and structural variation, we evaluated whether TE-associated variants displayed distinct patterns of differentiation compared to non-TE-associated mutations.

To evaluate whether chromosomal rearrangements in Mayotte *D. yakuba* are more likely to reflect standing variation or new mutations, we compared their allele frequency patterns with their transposable element (TE) associations. Newly arising mutations were defined as rearrangements with a nonzero allele frequency in Mayotte but an allele frequency of zero in the mainland population. We grouped significantly differentiated rearrangements into three categories based on TE involvement: (1) no TE association, (2) associated with ectopic recombination between TE copies, and (3) associated with TE insertions near rearrangement breakpoints.

**Figure 3:**
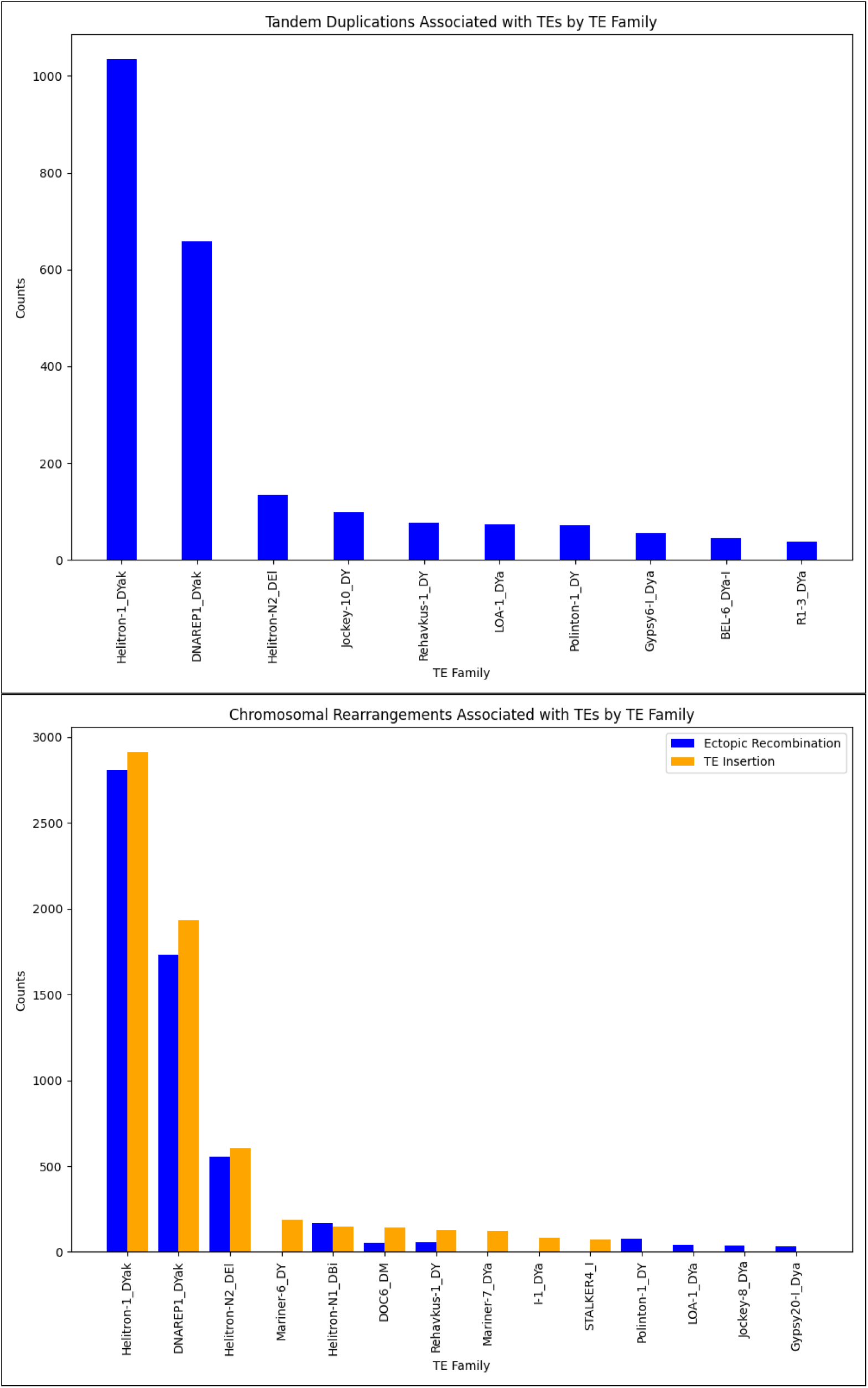
TE family associations for structural variants (SVs) in the Mayotte population.(A) Tandem duplications associated with nearby TEs, shown by TE family.(B) Chromosomal rearrangements associated with TEs, separated by inferred mutation mechanism: ectopic recombination (blue) or TE insertion (orange). Only TE families with ¿10 associated events in at least one category are shown.

For chromosomal rearrangements, TE-associated variants were significantly enriched among newly arising mutations (38/51; *χ*^2^ = 51.73, *p <* 3.50×10*^−^*^3^), whereas ectopic recombination events were primarily observed in rearrangements derived from standing variation (132/151; *χ*^2^ = 13.874, *p <* 5.20 × 10*^−^*^17^). These results indicate that TE insertions are frequently observed alongside new structural rearrangements, while ectopic recombination appears more associated with pre-existing variation.

For tandem duplications, we asked whether TE proximity differs between newly arising duplications and those derived from standing variation. Tandem duplications can arise through a variety of mutational processes, including ectopic recombination between repeated sequences, replication slippage, non-homologous end joining (NHEJ), and gene conversion, many of which do not involve TEs directly [45–47]. While some duplications show evidence of TE-mediated ectopic recombination at their boundaries, distinguishing specific mechanisms is challenging without high-resolution breakpoint data. As such, we categorized tandem duplications as either TE-associated or non-TE-associated, based on the presence of nearby TEs, to broadly capture potential TE influence. Chi-square analysis revealed no significant difference in TE associations between newly arising and standing duplications (*χ*^2^ = 0.27, *p* = 0.3). This suggests that TE-mediated dynamics may play a less pronounced role in early duplication formation compared to rearrangements.

The absence of TE enrichment among newly arising duplications suggests that TE-mediated processes may play a more limited role in early tandem duplication formation. Instead, duplications may initially arise through other mechanisms, such as replication slippage or NHEJ, with TE-associated ectopic recombination contributing to structural refinement or expansion over time [46]. This pattern raises the possibility that TE involvement in tandem duplication dynamics becomes more apparent over longer evolutionary timescales.

**Figure 4:**
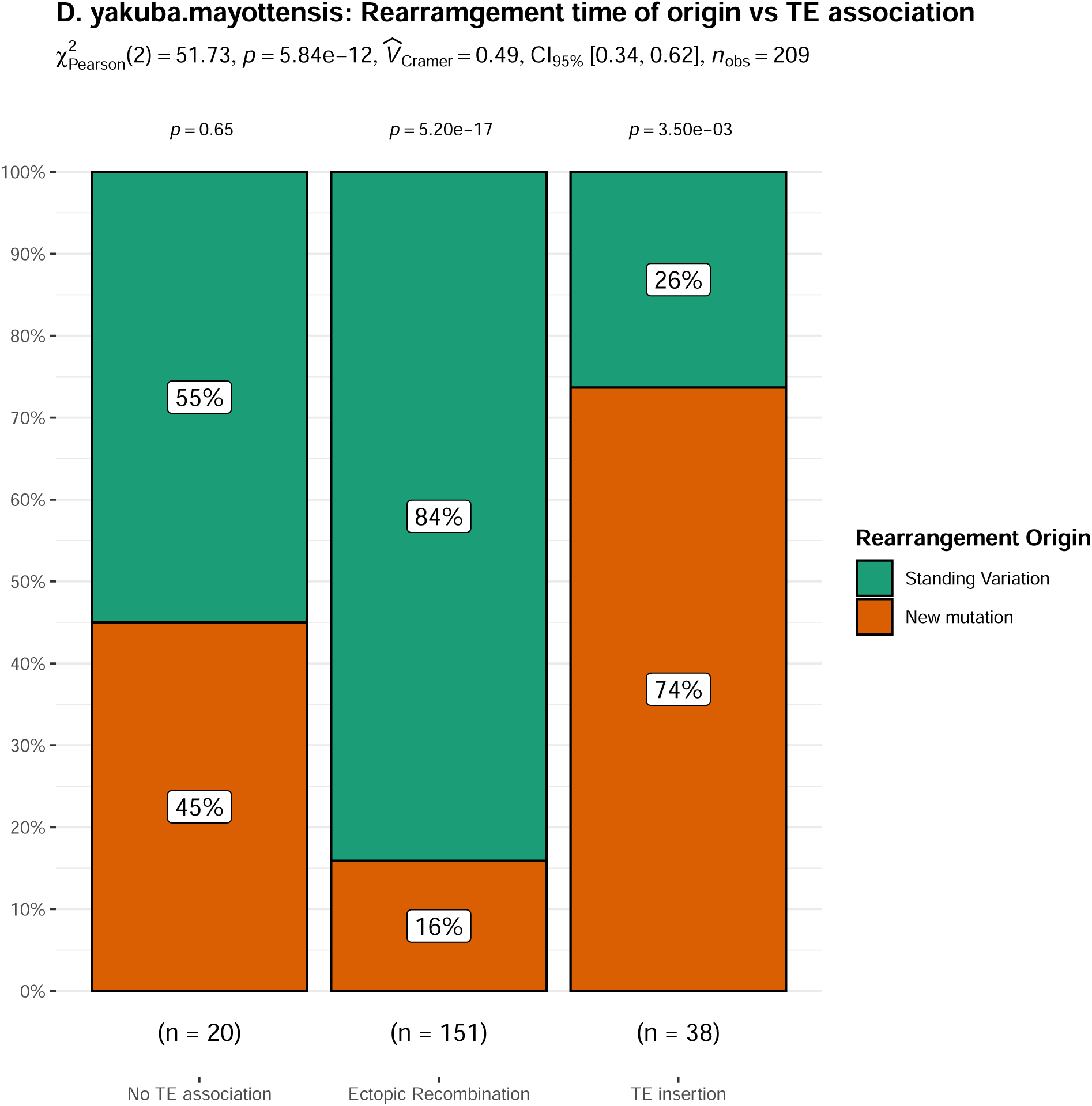
Bar plots showing the proportions of significantly differentiated rearrangements associated with TE insertions (left) and TE-mediated ectopic recombination (right) among newly arising variants (blue) and those derived from standing variation (orange). Rearrangements associated with TE insertions are significantly enriched among newly arising mutations (*χ*^2^ = 51.73, *p <* 3.50 × 10*^−^*^3^), suggesting that TE insertions are a major driver of novel structural variants shortly after colonization. In contrast, rearrangements linked to ectopic recombination are significantly more likely to be derived from standing variation (*χ*^2^ = 13.874, *p <* 5.20 × 10*^−^*^17^). The presence of these differentiated standing variants suggests that selection may have acted to retain or elevate their frequency following colonization. Together, these results indicate that different TE-mediated mechanisms contribute to chromosomal rearrangements over different evolutionary timescales, with TE insertions generating new variants and ectopic recombination shaping pre-existing variation potentially favored by selection.

Coding SNPs were also assessed for feature association by comparing the proportions of synonymous and nonsynonymous mutations among significantly differentiated variants. This analysis revealed a highly significant enrichment of nonsynonymous SNPs (*χ*^2^ = 164.57, *p <* 1.14 × 10*^−^*^37^). Among the SNPs showing significant allele frequency shifts between Mayotte and mainland populations, a larger proportion resulted in amino acid changes compared to synonymous substitutions, consistent with broader patterns in which mutation class influences differentiation.

### 3.4 Functional analysis of genes in island populations

Functional annotation analysis provides insight into enriched gene functions associated with identified genetic variation. Identifying functional enrichment may reveal biological pathways contributing to local adaptation. Comparing functional enrichment across different mutation classes, chromosomal rearrangements, tandem duplications, and coding SNPs, can help determine whether genetic novelty is mutation-limited in these structural and sequence-level variants.

In Mayotte *D. yakuba*, we identified 785 genes overlapping or within regulatory distance (+/-5kb) of all identified chromosomal rearrangements. Among these, 38 genes were associated specifically with significantly differentiated rearrangements. These genes clustered into 18 functional groups, but no gene ontology (GO) categories were significantly enriched, either when considering all rearrangements or only significantly differentiated ones. This suggests that chromosomal rearrangements impact a broad range of biological functions, but their effects are distributed across multiple biological processes rather than concentrated in specific functional categories. While individual rearrangements may have important functional consequences, they do not appear to disproportionately affect any single enriched pathway. For tandem duplications, we identified 235 genes associated with significantly differentiated duplications in the Mayotte population. These genes clustered into 7 functional groups, with a single enriched category related to calcium-dependent carbohydrate-recognition domains (C-lectins) (enrichment score, *ES* = 3.65).

**Figure 5:**
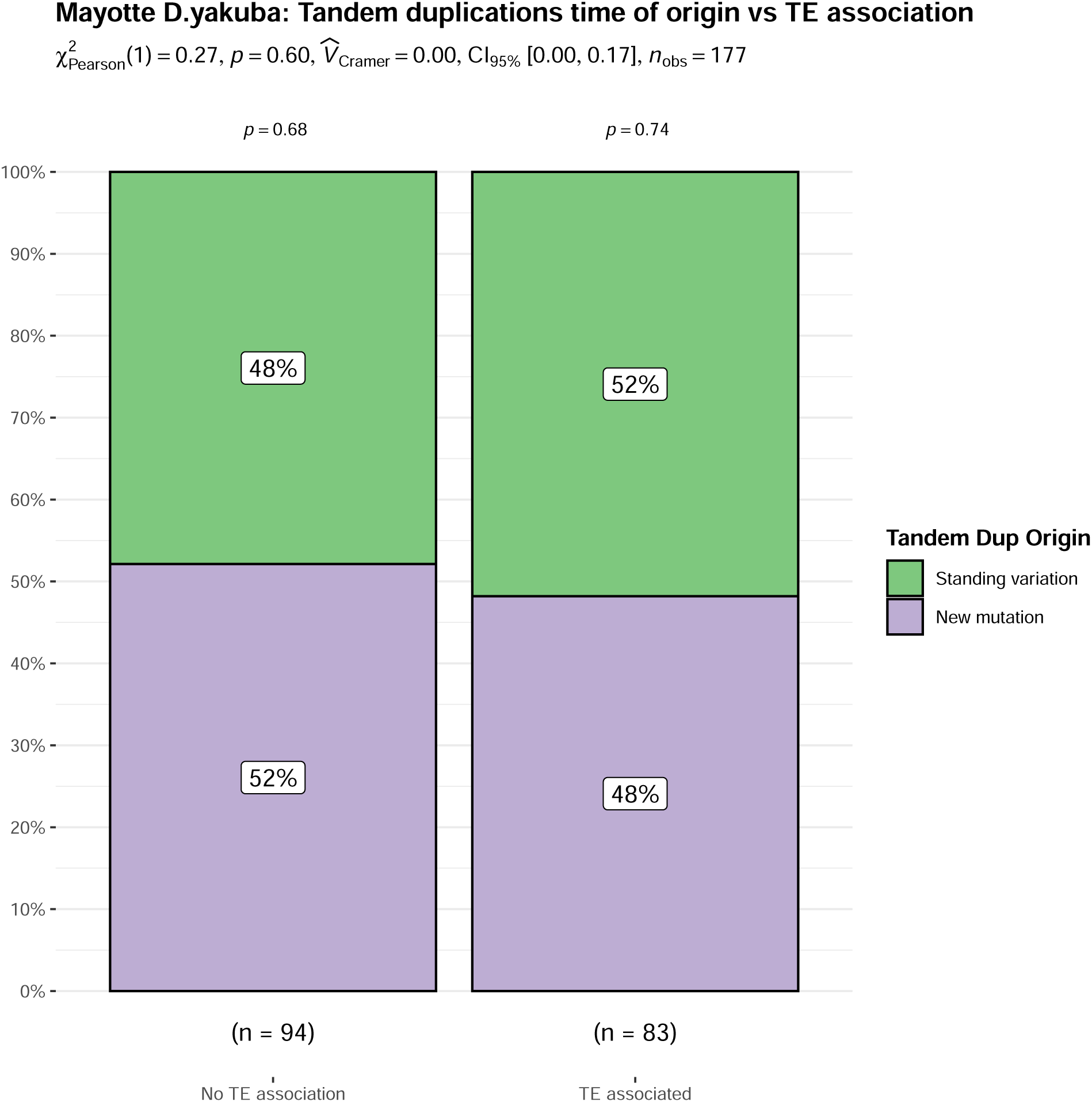
Bar plot showing the proportion of TE-associated versus non-TE-associated tandem duplications among newly arising duplications (green) and those derived from standing variation (purple). Chi-square analysis revealed no significant difference in TE association between these two classes (*χ*^2^ = 0.27, *p* = 0.3), indicating that TE-mediated tandem duplication events do not preferentially occur in new versus standing variants. This contrasts with chromosomal rearrangements, where TE associations are significantly enriched in newly arising mutations (see Figure 6). These results suggest that TE influence on tandem duplications may be more limited during early divergence and could become more relevant over longer evolutionary timescales.

**Figure 6:**
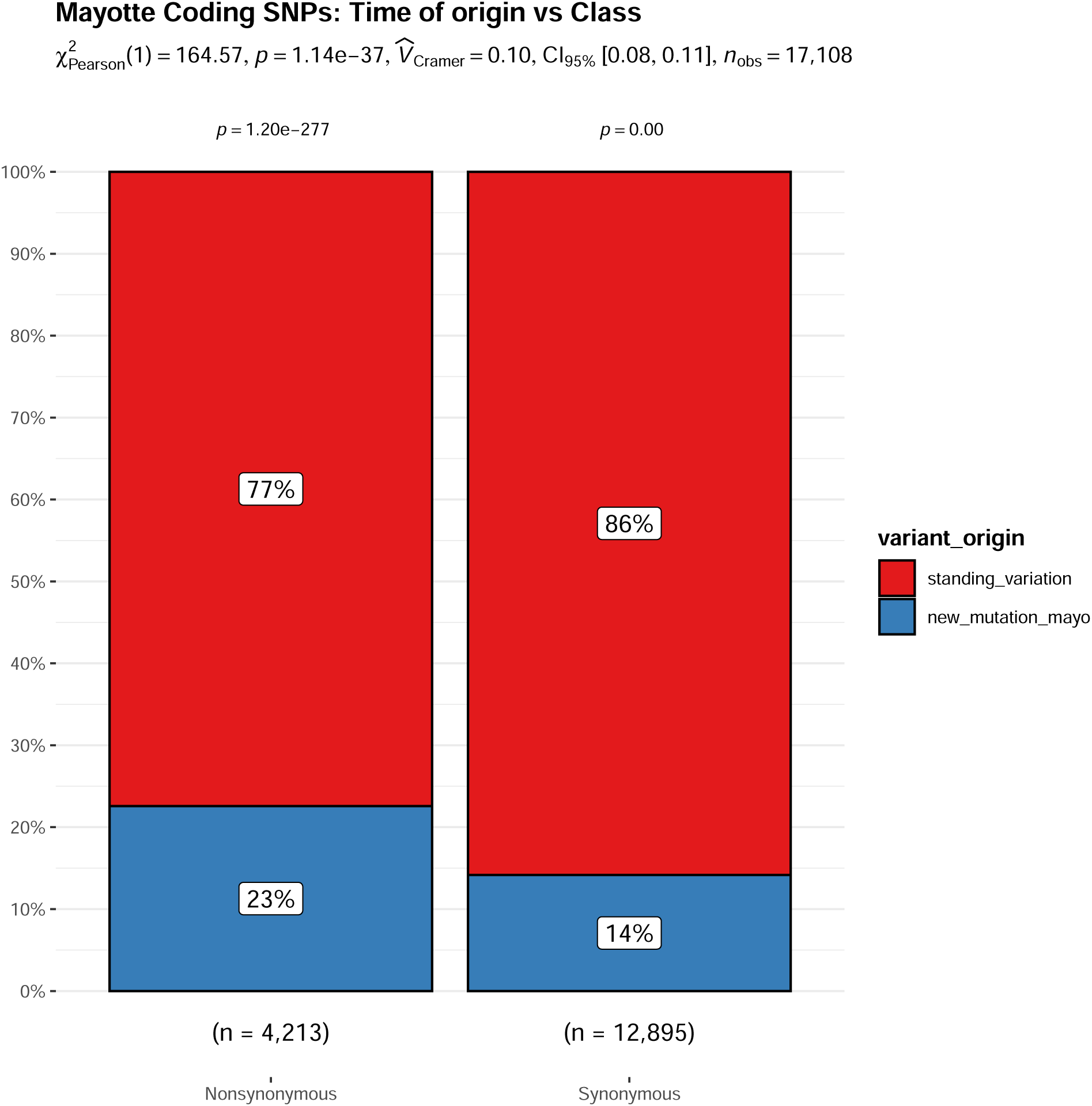
Bar plots showing the proportions of synonymous and nonsynonymous coding SNPs with significant differentiation statistics between the Mayotte *D. yakuba* and mainland population. The proportion of SNPs that are novel to the Mayotte population are shown in blue, and standing variants, SNPs seen in both the island and ancestral population, are shown in red. Chi-square analysis revealed a significant enrichment of nonsynonymous SNPs among differentiated sites (*χ*^2^ = 164.57, *p <* 1.14 × 10*^−^*^37^), indicating that a higher proportion of allele frequency shifts between Mayotte and mainland populations result in amino acid changes compared to synonymous substitutions, consistent with broader patterns of mutational class influencing genomic differentiation.

RepeatMasker analysis of the *D. yakuba* reference genome identified multiple Class 2 DNA transposons, including piggyBac, Transib, and Helitron elements, near these C-lectin tandem duplications. These transposons, particularly those known to promote ectopic recombination, suggest that TE activity may have facilitated copy number expansion at this locus.

C-lectins are involved in carbohydrate binding and immune response, but their specific functional role in these populations remains unclear. While this enrichment could suggest a role in local adaptation, determining whether these duplications are functionally relevant would require additional gene expression analysis.

### 3.5 Convergence and parallelism through tandem duplications in island populations

To investigate whether parallel genetic changes have occurred in isolated island populations, we analyzed tandem duplications shared between Mayotte *D. yakuba* and the mainland population. Specifically, we examined whether particular gene regions exhibited an excess of duplications in Mayotte, which could indicate convergent evolution driven by similar ecological pressures.

In Mayotte *D. yakuba*, we identified a hotspot of tandem duplications near the calciumdependent carbohydrate-recognition domain (C-lectin) gene region. This region contains 4 tandem duplications in the mainland population and 13 in Mayotte *D. yakuba*. Notably, 3 of the Mayotte duplications exhibit significant differentiation from the mainland population, suggesting that certain duplications have increased in frequency following island colonization. Conversely, three duplication calls exhibit moderate allele frequency in both Mayotte and the mainland, suggesting that some C-lectin duplications may have existed as low-frequency standing variation before becoming more prevalent in the island population. This pattern provides evidence that both standing variation and new mutations contribute to the genetic landscape of C-lectin tandem duplications.

The enrichment of C-lectin domains in tandem duplications is significant across all duplications in Mayotte. However, this enrichment diminishes when considering only significantly differentiated duplications, suggesting that the overall pattern is driven more by mutational dynamics than by allele frequency shifts indicative of selection. For this specific case, this may indicate that mutation, rather than strong selection, is the primary driver of duplication formation. Nonetheless, a subset of these duplications appears highly diverged, highlighting the interplay between mutation and selection in shaping genetic diversity in island populations.

## 4 Discussion

### 4.1 Comparing Divergence on Short vs. Long Evolutionary Timescales

On short evolutionary timescales, the emergence and retention of genetic variation is shaped by distinct mutational mechanisms and demographic processes. Single nucleotide polymorphisms (SNPs) accumulate steadily through relatively stable mutation rates [10,28], whereas structural variants (SVs), including chromosomal rearrangements and tandem duplications, can arise via more dynamic mutational pathways, particularly those facilitated by transposable elements (TEs) [12, 48]. Some SVs reflect standing genetic variation inherited from ancestral populations, while others represent newly arisen mutations [2, 4, 12].

Our results from Mayotte *D. yakuba* highlight how genomic differentiation progresses shortly after colonization (∼10,000 - 28,000 years ago), allowing us to assess the relative roles of SNPs, tandem duplications, and rearrangements early in divergence. To contextualize these findings, we compared our chromosomal rearrangement results with previously studied patterns of rearrangement differentiation in two island populations from São Tomè: the recently colonized São Tomè *D. yakuba* (∼10,000 years) and the longer-isolated *D. santomea* (∼500,000 years) [16]. This comparison provides insight into how the relative importance of chromosomal rearrangements in divergence may shift with increasing isolation time and ecological differentiation [15, 49].

Together, these comparisons suggest a general pattern: adaptive changes in Mayotte *D. yakuba* are partially constrained by mutation for SNPs, tandem duplications, and rearrangements formed via ectopic recombination, as the variants under selection are largely drawn from standing genetic variation. In contrast, TE insertions and TE-mediated rearrangements appear less constrained, arising more readily as new mutations and providing a mechanism by which novel adaptive variation can emerge even on short timescales [15, 49]. This pattern may also reflect a stronger impact of Haldane’s sieve on structural variants [36]. Because SVs often have large effects and arise at low frequencies, newly occurring variants are more likely to be lost to purifying selection before reaching appreciable frequency [1, 50, 51].

### 4.2 Early TE Activity and Structural Variant Formation in Island *D. yakuba*

Transposable elements (TEs) are well known drivers of structural genome evolution, frequently giving rise to mutations such as chromosomal rearrangements and tandem duplications. These SVs differ in scale and mechanism where rearrangements typically involve large scale genome restructuring, whereas tandem duplications alter copy number at more localized regions [48]. TEs can influence the formation of both classes through distinct mutational pathways, such as insertions or facilitating ectopic recombination [11, 12]. In recently colonized *D. yakuba* populations like those on Mayotte and São Tomè, TEs may contribute to early genomic divergence, but the degree to which these variants persist and influence adaptation remains unclear.

Our results suggest that TEs are disproportionately associated with chromosomal rearrangements in Mayotte *D. yakuba* genome wide. Chromosomal rearrangements (57.9%) and tandem duplications (35.2%) are frequently associated with nearby TEs, substantially higher than expected given that TEs comprise approximately 12% of the *D. yakuba* genome [32]. These findings reinforce the view that TE activity plays a central role in generating structural variation, some of which fortuitously become targets of selection in recently colonized island populations [16, 32, 52].

Our analyses show TE insertions disproportionately characterize significantly differentiated chromosomal rearrangements as new mutations in Mayotte *D. yakuba*, despite a relatively short period of isolation. This pattern aligns with observations from *D. santomea*, which similarly exhibits a high proportion of diverged rearrangements associated with new TE insertions [16]. In contrast, the proportion of rearrangements from standing variation in Mayotte more closely matches that in São Tomè *D. yakuba*, populations with more comparable colonization timelines. This suggests that while standing variation serves as the immediate reservoir of genetic diversity following colonization, new TE insertions can provide an early and persistent source of novel mutations that may convey selective advantages. We see that TE insertions, which are disproportionately new mutations, play a surprisingly prominent role in early divergence, a pattern that contrasts with expectations based on SNP dynamics where new mutations typically arise more slowly [1, 2, 10]. Beyond direct insertions, TEs can mediate ectopic recombination between dispersed copies, giving rise to complex rearrangements. While diverged TE insertions seem to contribute to divergence early, rearrangements from TE-facilitated ectopic recombination show an equally striking skew towards standing variation This difference in mutational mechanisms and variant origins highlights how structural variation evolves through multiple pathways.

### 4.3 Delayed Contribution of TE-Mediated Ectopic Recombination to Divergence

Although transposable elements (TEs) can provide abundant raw material for structural genome evolution, our results suggest that ectopic recombination between TEs plays a limited role in generating novel chromosomal rearrangements on short evolutionary timescales. [11, 12]. In contrast to findings seen in *D. santomea*, where TE-facilitated ectopic recombination is seen frequently among diverged rearrangements, novel TE-driven ectopic rearrangements are largely absent in both the São Tomè and Mayotte *D. yakuba* populations.

Novel ectopic recombination-driven rearrangements are rare among the Mayotte *D. yakuba*, a result mirrored among the São Tomè *D. yakuba* population. This means rearrangements associated with ectopic recombination are primarily derived from standing variation among the *D. yakuba* populations with more recent colonization timelines. This suggests that rather than generating entirely new rearrangements that rapidly diverge, ectopic recombination-associated variants were already present at low frequencies in the ancestral population and subsequently increased in frequency in the island populations [2]. If recombination between dispersed TE copies frequently produces large or disruptive rearrangements, purifying selection may remove these newly arising variants before they reach detectable frequencies [15]. This filtering effect is consistent with Haldane’s sieve [50], which predicts that beneficial variants arising as new mutations are less likely to fix if they are recessive or deleterious when rare. In contrast, pre-existing ectopic recombination-associated rearrangements may have already undergone some degree of filtering in the ancestral population, allowing these pre-tested mutations to persist and contribute to differentiation between island and mainland populations.

This pattern is supported when comparing this pattern to that of *D. santomea*, a species that has been isolated for approximately 500,000 years and exhibits a much stronger association between ectopic recombination and significantly differentiated rearrangements [16]. These results suggest that while ectopic recombination is largely absent from novel structural variants on short timescales, it may become more prominent in shaping divergence over longer evolutionary periods [53]. One possible explanation is that selection filters which rearrangements persist, allowing ectopic recombination-driven SVs to accumulate over time [15]. Ultimately, our findings indicate that TE facilitated ectopic recombination may be an important driver of divergence, but primarily through pre-existing structural variation rather than generating novel mutations on short timescales. While TE insertions provide an immediate source of genetic novelty, recombination driven structural variants may require time to generate new mutational substrate before playing a meaningful role in genome evolution. Given these mutational dynamics, a key question is whether structural variants observed in both Mayotte and São Tomè *D. yakuba* represent independent parallel evolution or are instead shaped by shared ancestral variation and recurrent mutational processes [37, 39].

### 4.4 Functional Dynamics of Diverged Mutations

In Mayotte *D. yakuba*, significantly differentiated SNPs were widely distributed across the genome but did not show enrichment for any specific functional categories. This pattern suggests polygenic adaptation, where subtle shifts across many loci contribute to early divergence without clustering in known biological pathways [54]. Tandem duplications, by contrast, showed focused enrichment for calcium-dependent carbohydrate-recognition domains (C-lectins), pointing to the possibility of gene family expansion or localized selection targeting immune-related or environmental response functions.

Chromosomal rearrangements, while overlapping many genes, did not show functional enrichment at the pathway level. This is consistent with earlier findings in São Tomè *D. yakuba* [16], where rearrangements affected a broad array of gene functions but lacked concentrated enrichment. In contrast, *D. santomea*, which has been isolated longer, showed functional enrichment among significantly differentiated and differentially expressed rearrangements, particularly in stress-response pathways [16]. This suggests that the adaptive relevance of rearrangements may take longer to emerge, potentially requiring extended periods of selection.

These comparisons imply that different mutation classes contribute to functional divergence in distinct ways. SNPs and duplications may facilitate more rapid shifts in gene regulation or dosage, while rearrangements exert broader, more diffuse effects that are slower to align with clear functional outcomes. Indeed, several pronounced SNP differentiation peaks in Mayotte *D. yakuba* overlap known inversion breakpoints, including a prominent X-linked inversion [41, 43]. These peaks coincide with elevated transposable element (TE) activity, including 372 insertions from the SAR2-DM element family. Such TE-rich inversion regions may preserve adaptive combinations of mutations and contribute to local genomic instability, compounding their role in divergence [41, 43, 55].

Finally, the observation that several functionally relevant duplications are associated with nearby TEs suggests that TE activity may promote localized functional innovation. In particular, TE-mediated mechanisms such as ectopic recombination or replication slippage may have facilitated the origin and spread of the C-lectin duplications, linking mutational dynamics to early functional divergence in island populations.

### 5.5 Shared Structural Variation and the Potential for Convergent Evolution

We identified several chromosomal rearrangements in Mayotte *D. yakuba* that overlap with variants previously reported in *D. yakuba* residing on São Tomè. These overlaps often occur in regions of high transposable element (TE) density or near known rearrangement breakpoints, suggesting that certain genomic regions may be structurally constrained and prone to recurrent structural change [11, 12]. In some cases, the rearrangements are structurally similar across populations, raising the possibility that these regions experience repeated mutation or retain ancestral variants across both island lineages.

More broadly, our data show that structural variants, particularly those associated with TE activity, tend to recur in specific regions of the genome. This pattern is consistent with mutational bias, where certain loci are more prone to rearrangement due to sequence composition, repeat content, or local structural instability [37, 56]. In some cases, variants arising in these mutation-prone regions may confer fitness advantages and increase in frequency, especially if they affect traits relevant to novel selective pressures. Both mutational constraint and selection likely contribute to the repeated appearance and persistence of structural variants in similar genomic regions across populations.

This has important implications for interpreting repeated genetic patterns. In systems with strong mutational biases, parallel genomic changes may arise even in the absence of identical selective pressures [37, 39]. Conversely, similar ecological conditions on different islands could still favor some of these recurrent variants if they happen to confer fitness benefits. These scenarios are not mutually exclusive and can be difficult to disentangle without functional validation.

Our results also speak to classic ideas in evolutionary theory, like Fisher’s model of adaptive walks [38]. This model describes adaptation as a stepwise process, where populations move toward a fitness peak by fixing beneficial mutations. Early in that process, larger-effect mutations can be helpful, but as the population gets closer to an optimum, smaller-effect changes are more likely to improve fitness [14]. In this context, structural variants, especially those driven by TE activity, might play a bigger role early on, when large changes can open up new trait combinations or help populations respond quickly to environmental shifts. Later on, SNPs and smaller-scale mutations may take over, fine-tuning traits that are already close to optimal.

However, if large effect TE induced mutations are in fact not available among standing variation but rather arise from new mutation, then the trajectory of adaptive walks may differ. Smaller effects could spread first from the immediately available standing variation and larger effects arising later after new mutations appear in natural populations. As such, further understanding of the timing of evolutionary change may fundamentally alter our understanding of how evolutionary processes proceed in nature. The fact that some structural changes recur in the same regions across populations also suggests that mutation bias influences which evolutionary paths are actually used. Together, this points to a mix of mutation type, demographic history, evolutionary timeline, and selective pressure shaping how populations move through evolutionary space.

## 5 Methods

### Whole genome sequencing

By studying isolated populations, like those of the *Drosophila* found on the islands of Mayotte and São Tomè, we can assess the tempo of evolution, and if certain genetic mechanisms can disproportionately result in rapid adaptive evolution. DNA was extracted from flies flash frozen in liquid nitrogen following QIAamp Mini Kit (Qiagen) protocol without using RNase A. The resulting DNA samples were quantified (Qubit dsDNA HS assay kit, ThermoFisher Scientific), assessed for quality (Nanodrop ND-2000; A260/A280*>*1.8), and stored at -20°C. Illumina TruSeq Nano DNA libraries were prepared manually following the manufacturer’s protocol (TruSeq Nano DNA, RevD; Illumina). Briefly, samples were normalized to 100ng DNA and sheared by sonication with Covaris M220 (microTUBE 50; Sage Science). The samples were end repaired, purified with Ampure XP beads (Agencourt; Beckman Coulter), adaptors adenylated, and Unique Dual Indices (Table) ligated. Adaptor enrichment was performed using eight cycles of PCR. Following Ampure XP bead cleanup, fragment sizes for all libraries were measured using Agilent Bioanalyzer 2100 (HS DNA Assay; Applied Biosystems). The libraries were diluted 1:10 000 and 1:20 000 and quantified in triplicate using the KAPA Library Quantification Kit (Kapa Biosystems). Equimolar samples were pooled and the libraries were size selected targeting 400-700bp range to remove adaptor monomers and dimers using Pippen Prep DNA Size Selection system (1.5% Agarose Gel Cassette #CDF1510; Sage Sciences). Library pools (24 samples per lane) were run on an llumina HiSeq 4000 platform using the 150bp paired end (PE) Cluster Kit.

We aligned short read sequences to the *D. yakuba* reference genome *Prin Dyak Tai*18*E*2 2.1 and *Wolbachia* endoparasite sequence NC 002978.6 using bwa aln (version 0.7.17) [57], and resolved paired end mappings using bwa sampe. We sorted alignments by position using samtools sort (version 1.17) [58]. Sequence depth on sorted bam files was calculated using samtools depth -aa.

### Identification of Structural Variants and TEs

We identified abnormally mapping read pairs on different chromosomes and long-spanning read pairs greater than 100kb apart as signals of putative chromosomal rearrangements. Mutations with greater than 3 read pairs supporting were included among the chromosomal rearrangements. Rearrangement calls were then clustered across samples using in-house python, assuming that rearrangements both rearrangement breakpoints (putative start and stop position of the rearrangement) were both within 325 base pairs of each other on. Once clustered, where the min and max position of rearrangement breakpoints were used to represent the cluster, we associate all strains in said cluster with these bounds.

To identify heterozygous regions (later used to correct allele frequencies), we used a Hidden Markov Model (HMM) to parse SNP heterozygosity for each strain. In *D. yakuba*, some strains had reference strain contamination, resulting in no heterozygous or homozygous non-reference SNPs. We used a three-state HMM for *D. yakuba* to identify reference contamination, inbred haplotypes, and non-inbred haplotypes. Transitions probabilities were set to 10*^−^*^10^. Emission probabilities were set as heterozygosity for inbred regions would be 0, and heterozygosity in non-inbred regions was *θ* = 0.01, with a lower threshold for probabilities on off-diagonals of *ɛ* = 0.00005 to prevent chilling effects of zero probability. Haplotype calls from the HMM were used to generate correct site frequency spectra for SNPs and structural variants given the variable number of chromosomes sampled across different parts of the genome.

Within inbred regions, we assume mutations are homozygous, and assign a genotype of 1 structural variant on 1 sampled chromosome. Within heterozygous haplotypes, read pair information alone cannot detect ploidy. For outbred regions with two chromosomes with different ancestry, we used coverage changes to distinguish hemizygous and homozygous mutations. We compared the average coverage of observed mutations to the average coverage across the entire chromosome in a strain looking for an observed coverage between 1.25 *<*= *x <*= 1.75 or 1.75 *< x <*= 2.25. Regions whose haplotypes were approximately 1.5x the average coverage of the chromosome are expected to be heterozygous, and regions who have approximately 2x the average are expected to be homozygous. False negative genotypes are possible when read pair support is insufficient to identify rearrangements de novo. For each mutation, if another strain showed 1.5x or 2x coverage changes and lesser support of 1 or 2 read pairs, allele frequencies were adjusted to avoid low frequency reads, similarly to prior work on gene duplications and rearrangements [18, 59].

Genotyping identifies mutations in populations that differ from the reference sequence, but on its own cannot identify which is ancestral and which is novel. To determine the ancestral state for rearrangements, we compared each variant to *D. teissieri*, we determine whether mutations are novel or if the ancestral state has been rearranged in the reference strain. To polarize the mutations we use BLASTn [60] at an E-value of 10*^−^*^20^ to compare the *D. yakuba* reference sequence for +/- 1kb of each rearrangement breakpoint to *D. teissieri* (*Prin_D_tei*_1_.1. We consider a mutation the ancestral state if both sequences map to the same location in *D. teissieri*, they have an alignment length greater than 150, share greater than 95 percent identity, and the two sequences overlap less than 10 percent in *D. teissieri*. Mutations where samples were identified as containing the ancestral state found in *D. teissieri*, allele frequencies from genotyping, *p*, were reversed to (1 − *p*) to reflect the reference as the new mutation. Comparing these frequencies between populations with an in house python script allows us to determine whether a mutation was new (found only in the mainland), or standing variation (found in both populations).

To match variants with transposable elements (TEs) we used BLAST to compare the variants to Repbase. Variants that mapped to TEs at an e-value of 10*^−^*^20^ were marked as TEs in our analysis. By using BLAST to compare both the origin and destination loci of the variant, we were able to see which variants mapped on either one or both sides of the rearrangement. (This was verified by checking the regions to make sure that they had unusual coverage compared to the average for that chromosome and strain.) Using BLAST, we compare each of the rearrangement breakpoints against the RepBase database [61].

### SNP Calling and Annotation Analysis

SNPs were identified by running bcftools mpileup (version 1.22) [58] and bcftools call (version 1.22) with the “-m” flag to create a VCF and corresponding index file for each sample. For SNP analysis, VCF files are filtered to contain only biallelic SNPs using bcftools view (version 1.22) with the “-m2 -M2 -v snps” flags.

Annotation was performed each VCF file using SNPGenie [62]. SNPGenie was run on each VCF which separated by chromosome to improve runtime, and on both the forward and reverse strand. The program requires a gene transcript file (GTF) to run, where the each gene reports one transcript per gene in the input GTF. To filter the GTF, we used AGAT-keep-longest-isoform (version 1.4.2) [63] to select only the longest transcript per gene for both the forward and reverse strands and create a filtered GFT for each strand.

SNPGenie reports information for each site, codon, transcript, and sample as a whole. For example, the total number of potential synonymous and nonsynonymous sites. Depending on the reported information, we merge the reported statistics of the forward and reverse strand according to literature. Output files are merged for all samples, to consolidate results.

### Population Branch Statistics and Pairwise Differentiation

Rearrangement frequencies were calculated by clustering across samples using an in-house python script. This script sorts all rearrangements by chromosome and position, placing them into bins with sizes determined by library insert size. Rearrangements in bins were clustered, and then compared to next door bins to avoid being missed as an edge case based on sorting. These frequencies are corrected residual heterozygosity created by inbreeding resistance at inversions similar to prior work [17], and are polarized based on the ancestral state of a rearrangement. Calculating the differences in the populations at rearrangements we can infer what changes have accumulated in the Mayotte *D. yakuba* since their separation from the mainland *D. yakuba*. With the allele frequency data for each variant and the separation by sub-population, we generate an SFS while accounting for uneven sample sizes across the genome. We projected allele frequencies in each subpopulation to a sample size of n=19 using a hypergeometric transformation, to evaluate the SFSs of populations with unequal population sizes [18, 64].

Differentiation between populations is a signature of local adaptation to changing environments. This differentiation can be measured through different means. This study benefits from a known ancestral mainland population that invaded new island environments that can be used as a population genetic ‘control’ to identify allele frequency changes. Simulations from other groups have shown that the difference in allele frequencies, Δ*p* can identify population differentiation during local adaptation. Δ*p*, and *F_ST_* contain overlapping information. Δ*p* contains directionality that *F_ST_* lacks, and may be more useful on moderate timescales when populations have had sufficient time for variants to spread in populations. It may identify population differentiation better than *F_ST_* while *F_ST_* may be more sensitive to changes while allele frequencies are low. We calculate Δ*p* and *F_ST_* identify structural variants that have spread in island environments.

Normalized PBS values were calculated using scikit-allel [65]. VCF files were loaded using the read-vcf method, and data were converted into genotype and allele count arrays required for the pbs function. PBS statistics were then computed for the mainland, Mayotte, and São Tomè *D. yakuba* populations.

By comparing the distribution to neutral SNPs, from bp 8-30 at first introns [66], in the genome we can determine the threshold of Δ*p* that should be used to determine if a mutation’s differentiation is beyond expectations for the allele frequency changes at neutral sites across the genome. These Δ*p* thresholds were calculated by chromosome using a Bonferroni corrected P-value of 0.05.

To determine whether mutations showed unusually high population differentiation we calculated a Bonferroni corrected 95% confidence interval, calculated by chromosome due to the distribution of Δ*p* varying by chromosome. Mutations that were greater than the upper bound of the C.I (2L,0.0936917; 2R,0.09405691; 3L,0.08218144; 3L,0.08330457; X,0.1462702) were considered significantly differentiated.

### Gene Ontology

Using the positions of the rearrangements we looked 5kb upstream and downstream of the mutations to see if their were genes either overlapping, or within regulatory distance. Variants that matched *D. yakuba* gene coordinates were mapped to their orthologs in *D. melanogaster* using Flybase [67]. Genes that had no orthologs were excluded. Using these orthologs we analyzed the functional gene annotations using DAVID [68] using the “low” stringency setting.

## Supporting information

AllSuppFigures

## 6 Acknowledgements

This work was supported by the National Institute of General Medical Sciences at the National Institutes of Health [R35-GM133376 to R.L.R] and the University of North Carolina, Charlotte [startup funding to R.L.R]; The authors thank Daniel Matute for collecting, maintaining, and sharing *Drosophila* stocks. We thank Cathy Moore for molecular sequencing assistance.

