## Supplementary material for "Genetic Novelty and Short-Term Evolutionary Dynamics in *Drosophila yakuba mayottensis*": AllSuppFigures

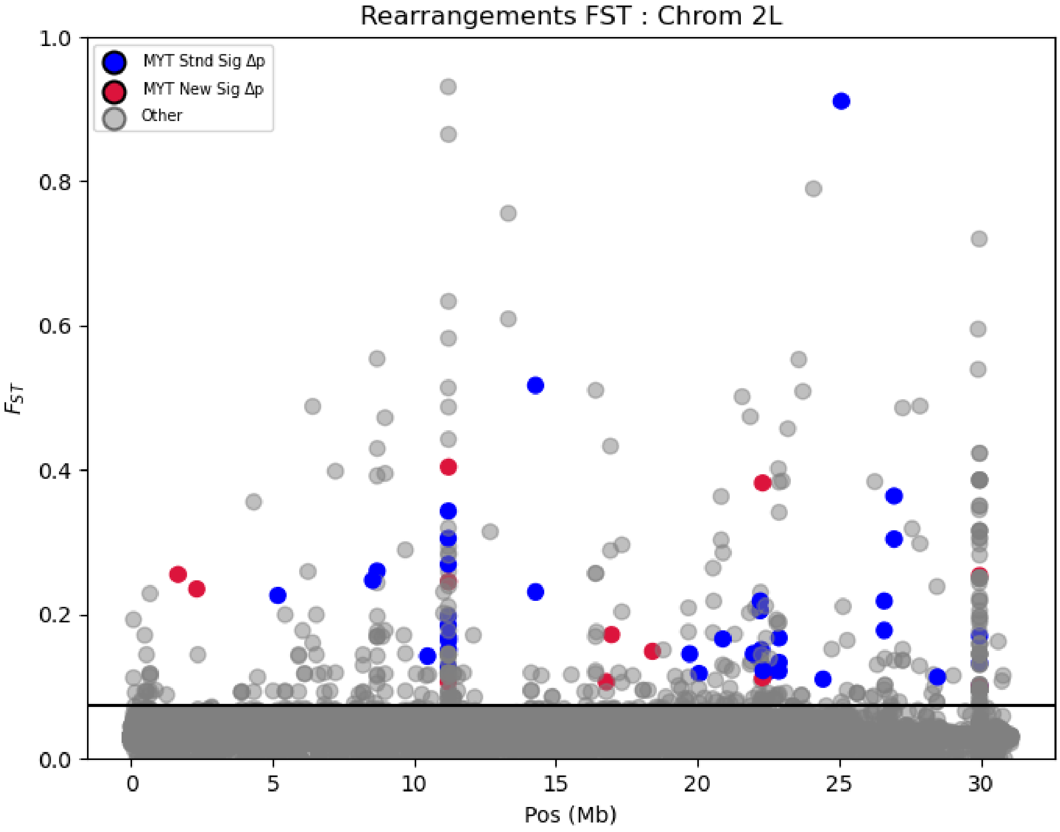


Figure S1: Multiple differentiation metrics (FST and $\Delta p$) for chromosomal rearrangements on the 2L chromosome. Plotting FST values for all rearrangements, and then highlighting those with significant allele frequency differences ($\Delta p$) for rearrangements disproportionately observed in the Mayotte *D. yakuba* population. Blue points represent significantly differentiated standing variation on Mayotte, and red points represent significantly differentiated new mutations on Mayotte. The horizontal black bar establishes the $\Delta p$ significance threshold.

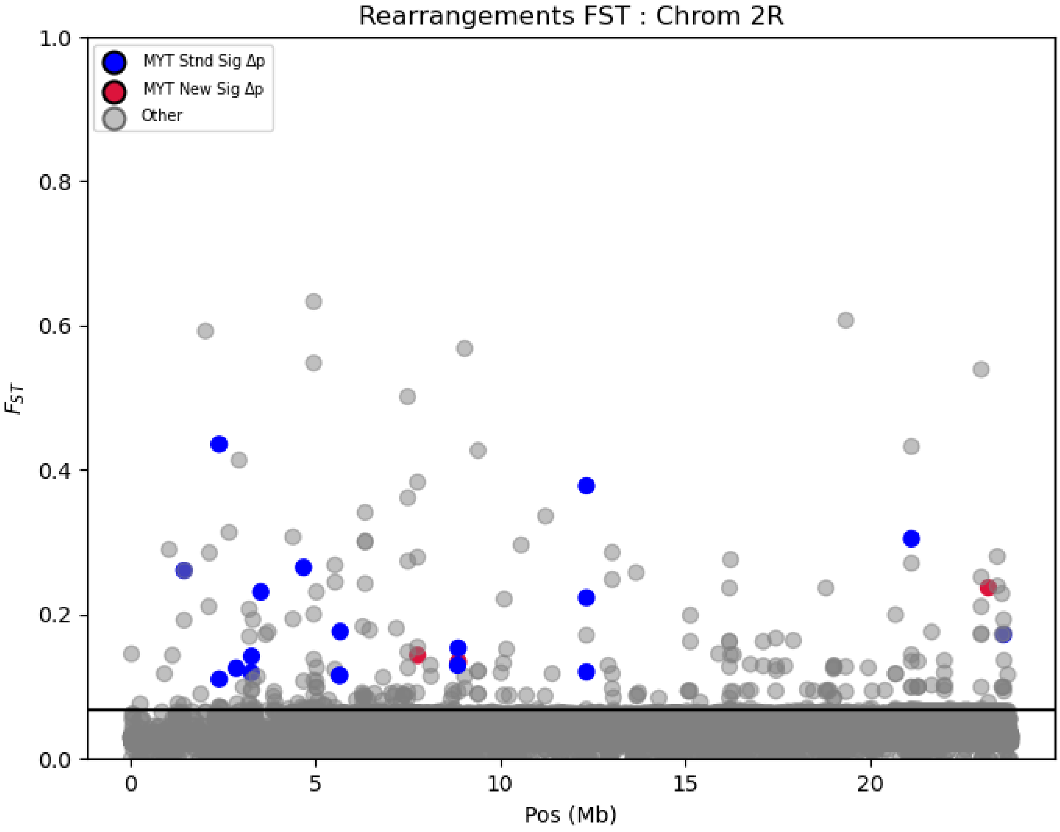


Figure S2: Multiple differentiation metrics (FST and $\Delta p$) for chromosomal rearrangements on the 2R chromosome. Plotting FST values for all rearrangements, and then highlighting those with significant allele frequency differences ($\Delta p$) for rearrangements disproportionately observed in the Mayotte *D. yakuba* population. Blue points represent significantly differentiated standing variation on Mayotte, and red points represent significantly differentiated new mutations on Mayotte. The horizontal black bar establishes the $\Delta p$ significance threshold.

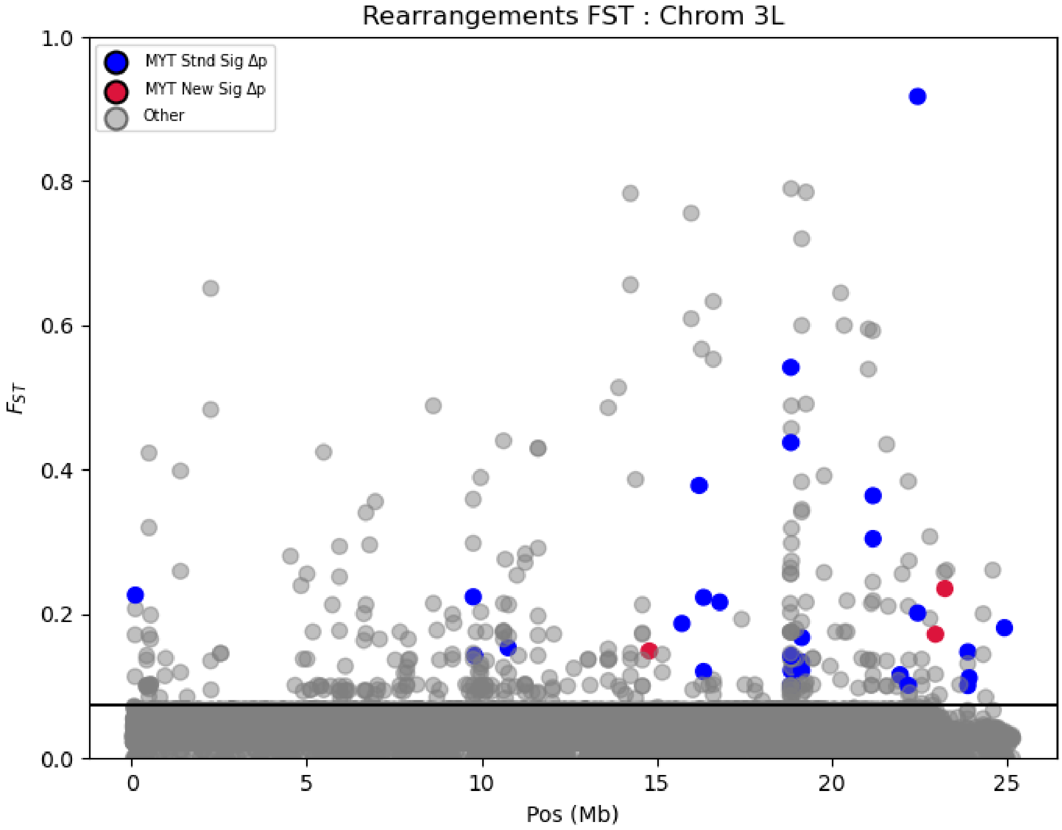


Figure S3: Multiple differentiation metrics (FST and $\Delta p$) for chromosomal rearrangements on the 3L chromosome. Plotting FST values for all rearrangements, and then highlighting those with significant allele frequency differences ($\Delta p$) for rearrangements disproportionately observed in the Mayotte *D. yakuba* population. Blue points represent significantly differentiated standing variation on Mayotte, and red points represent significantly differentiated new mutations on Mayotte. The horizontal black bar establishes the $\Delta p$ significance threshold.

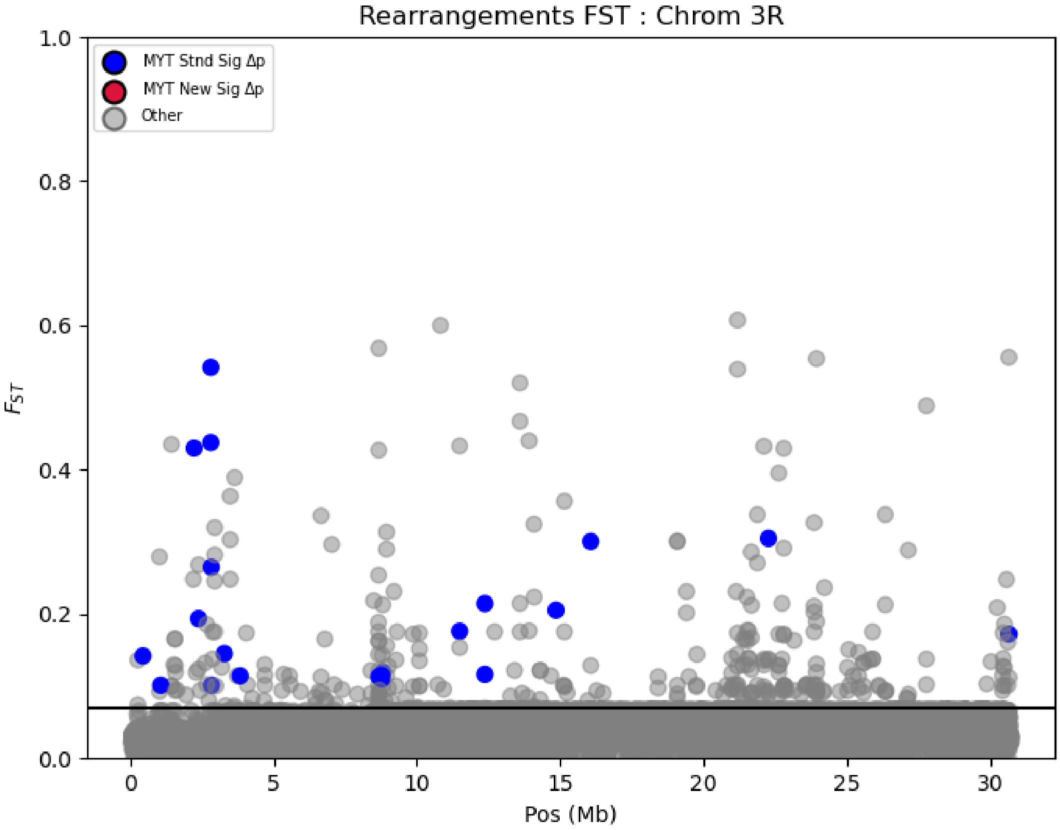


Figure S4: Multiple differentiation metrics (FST and $\Delta p$) for chromosomal rearrangements on the 3R chromosome. Plotting FST values for all rearrangements, and then highlighting those with significant allele frequency differences ($\Delta p$) for rearrangements disproportionately observed in the Mayotte *D. yakuba* population. Blue points represent significantly differentiated standing variation on Mayotte, and red points represent significantly differentiated new mutations on Mayotte. The horizontal black bar establishes the $\Delta p$ significance threshold.

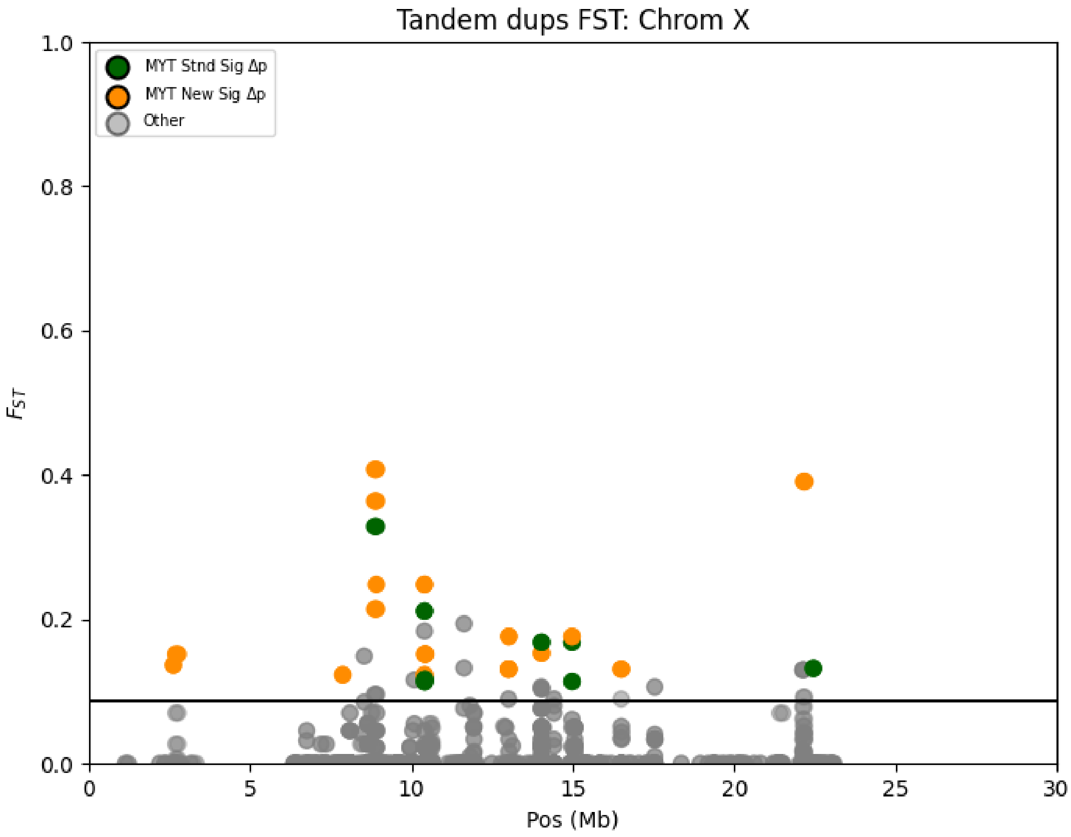


Figure S5: Multiple differentiation metrics (FST and $\Delta p$) for tandem duplications on the 2L chromosome. Plotting FST values for all duplications, and then highlighting those with significant allele frequency differences ($\Delta p$) for duplications disproportionately observed in the Mayotte *D. yakuba* population. Blue points represent significantly differentiated standing variation on Mayotte, and red points represent significantly differentiated new mutations on Mayotte. The horizontal black bar establishes the $\Delta p$ significance threshold.

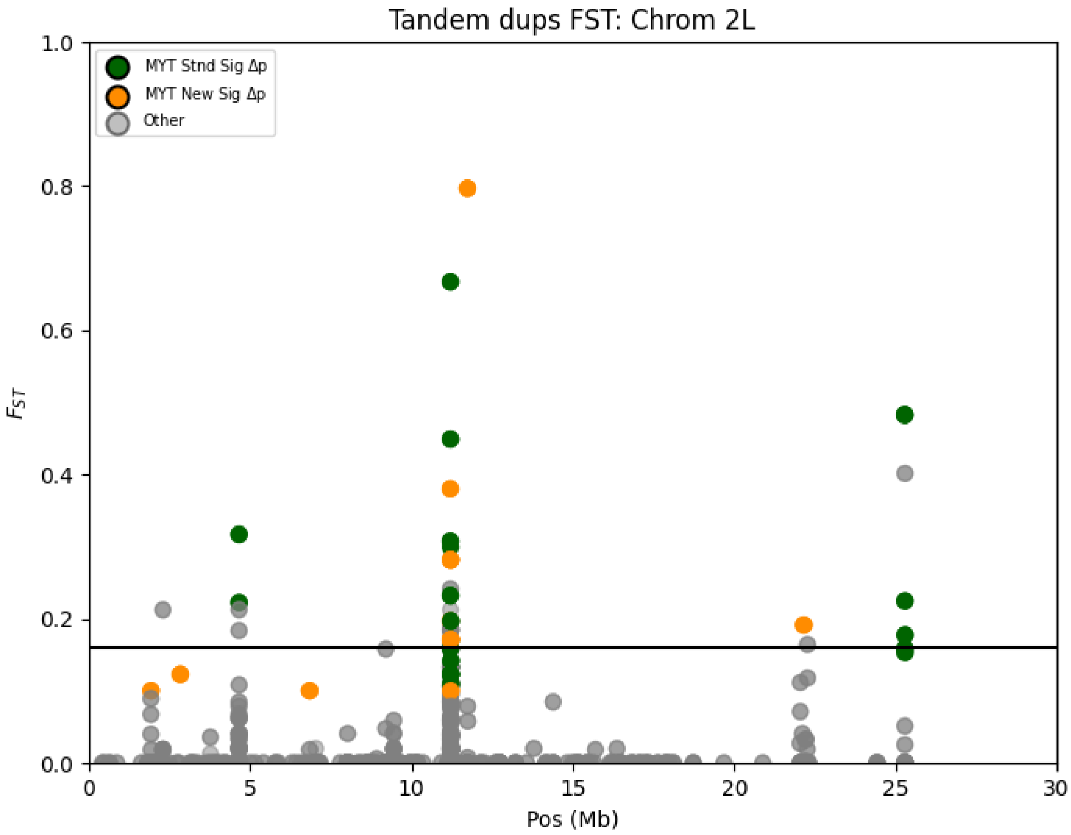


Figure S6: Multiple differentiation metrics (FST and $\Delta p$) for tandem duplications on the 2L chromosome. Plotting FST values for all duplications, and then highlighting those with significant allele frequency differences ($\Delta p$) for duplications disproportionately observed in the Mayotte *D. yakuba* population. Blue points represent significantly differentiated standing variation on Mayotte, and red points represent significantly differentiated new mutations on Mayotte. The horizontal black bar establishes the $\Delta p$ significance threshold.

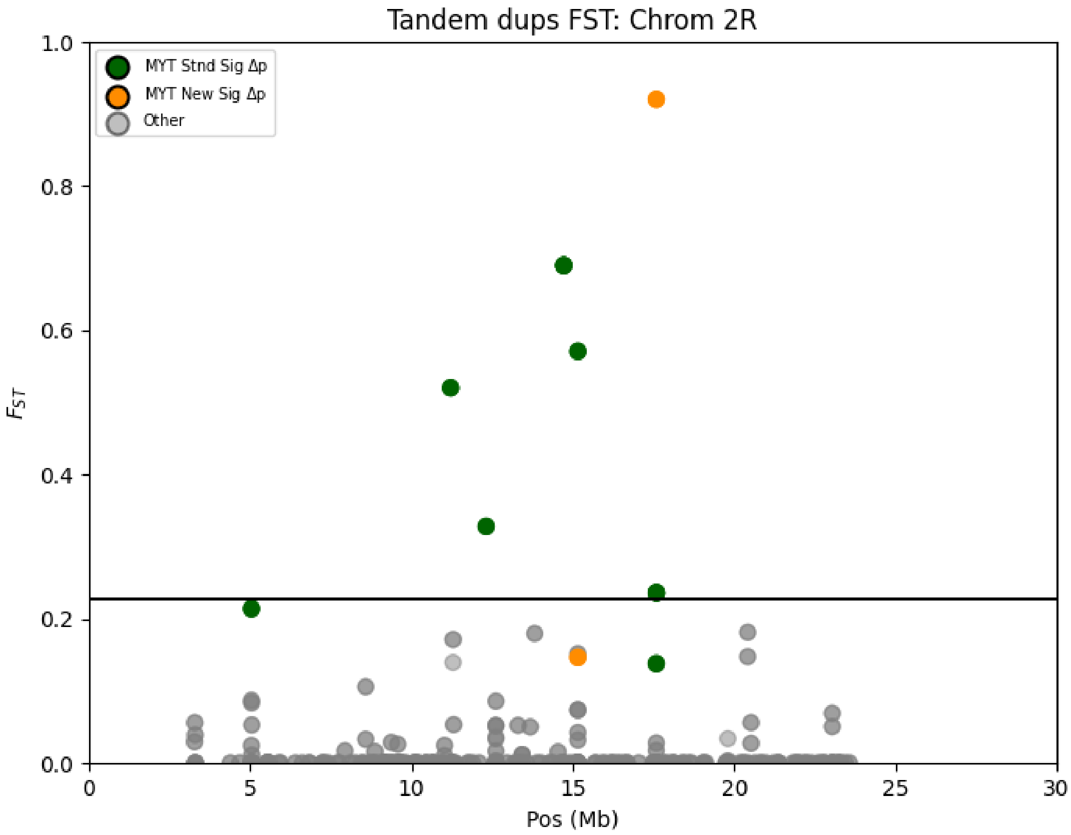


Figure S7: Multiple differentiation metrics (FST and $\Delta p$) for tandem duplications on the 2R chromosome. Plotting FST values for all duplications, and then highlighting those with significant allele frequency differences ($\Delta p$) for duplications disproportionately observed in the Mayotte *D. yakuba* population. Blue points represent significantly differentiated standing variation on Mayotte, and red points represent significantly differentiated new mutations on Mayotte. The horizontal black bar establishes the $\Delta p$ significance threshold.

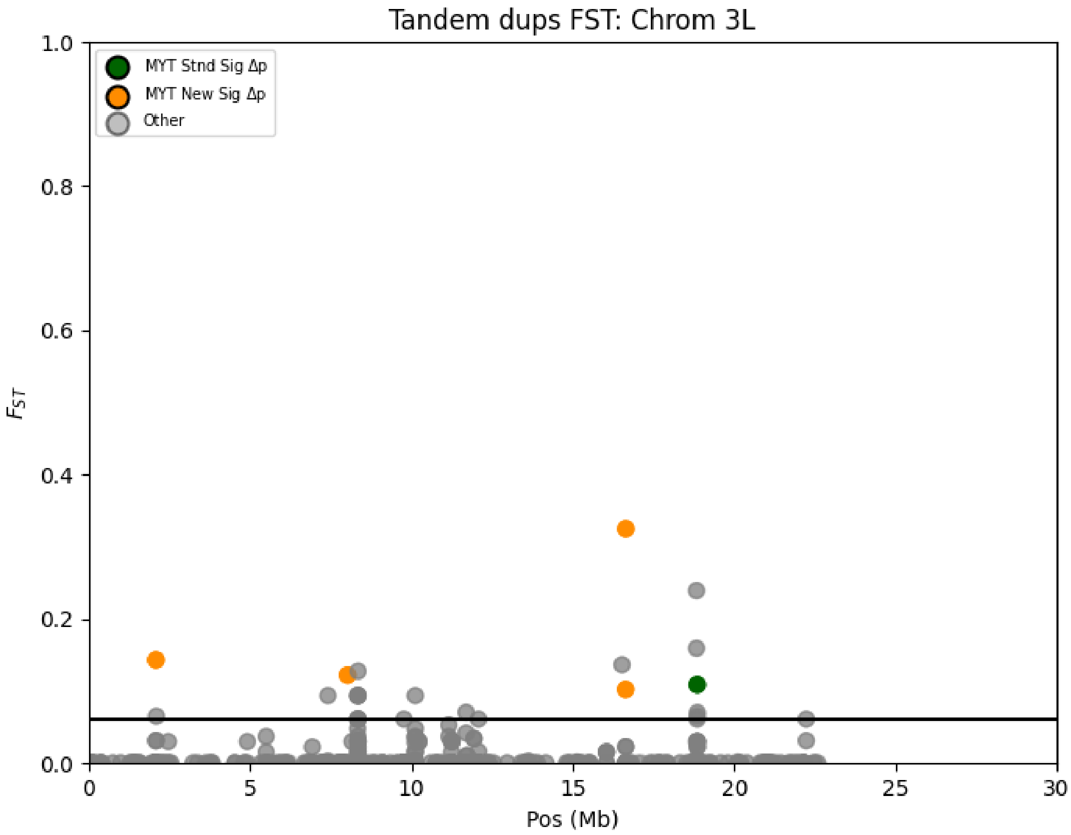


Figure S8: Multiple differentiation metrics (FST and $\Delta p$) for tandem duplications on the 3L chromosome. Plotting FST values for all duplications, and then highlighting those with significant allele frequency differences ($\Delta p$) for duplications disproportionately observed in the Mayotte *D. yakuba* population. Blue points represent significantly differentiated standing variation on Mayotte, and red points represent significantly differentiated new mutations on Mayotte. The horizontal black bar establishes the $\Delta p$ significance threshold.

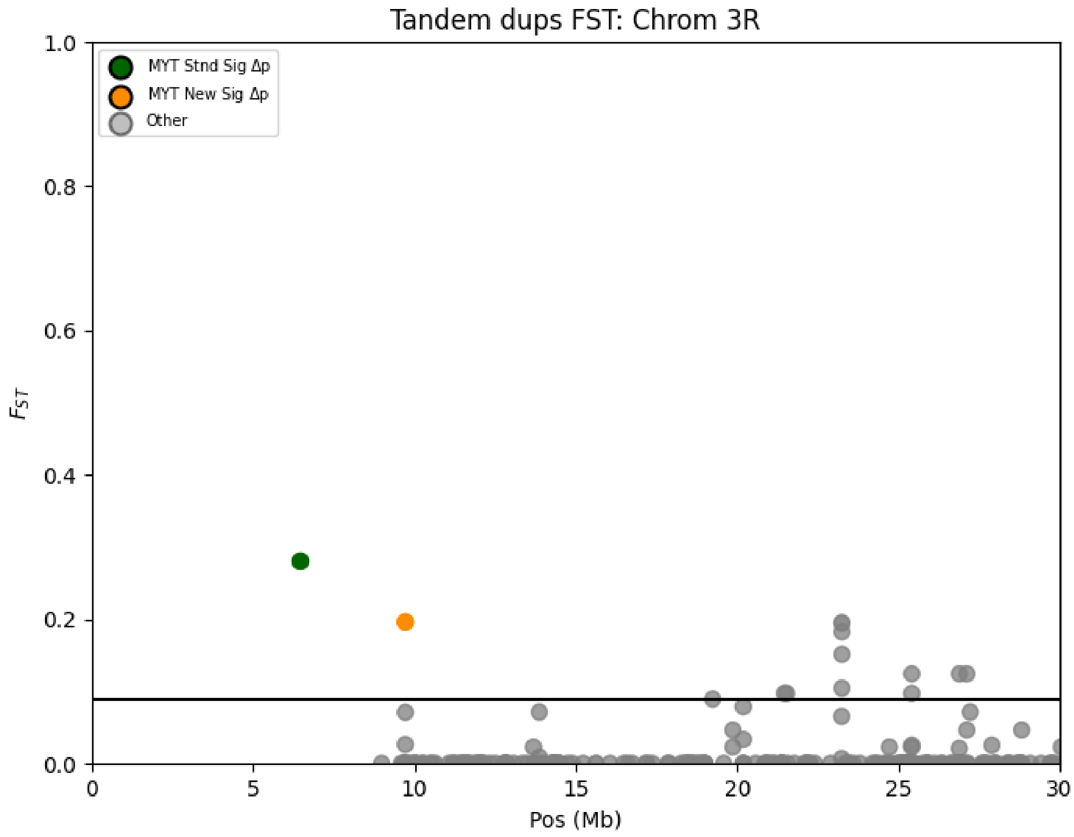


Figure S9: Multiple differentiation metrics (FST and $\Delta p$) for tandem duplications on the 3R chromosome. Plotting FST values for all duplications, and then highlighting those with significant allele frequency differences ($\Delta p$) for duplications disproportionately observed in the Mayotte *D. yakuba* population. Blue points represent significantly differentiated standing variation on Mayotte, and red points represent significantly differentiated new mutations on Mayotte. The horizontal black bar establishes the $\Delta p$ significance threshold.

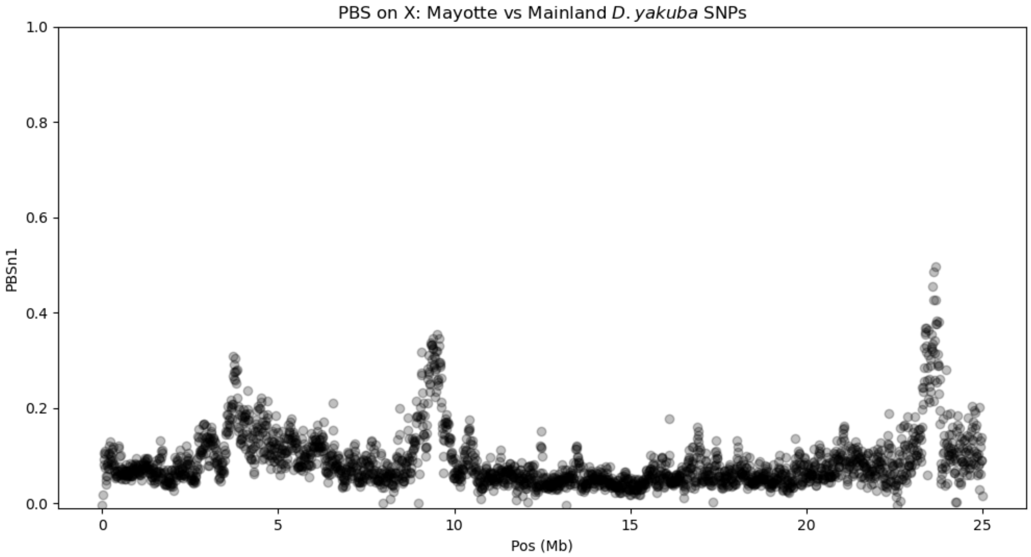


A

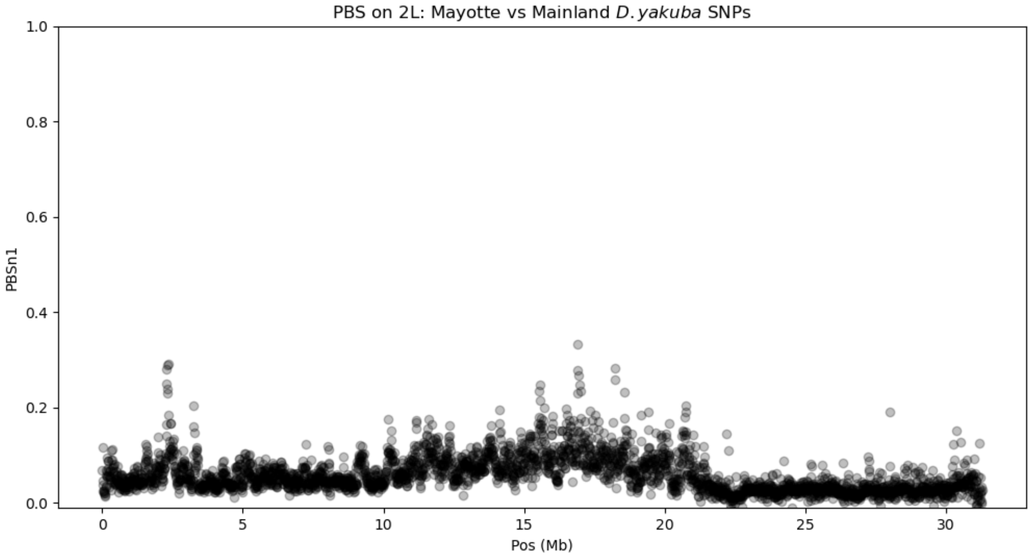


B

Figure S10: A) Site frequency spectrum of *D. santomea*. The increase in high frequency variants is much greater than that of the mainland *D. yakuba* B) Site frequency spectrum of the island *D. yakuba*. The increase in high frequency variants is slightly greater than that of the mainland *D. yakuba*, but not as much as *D. santomea*, likely related to divergence time C) Site frequency spectrum of mainland *D. yakuba* lacks the increase in high frequency variants that is present in both of the other sub-populations.

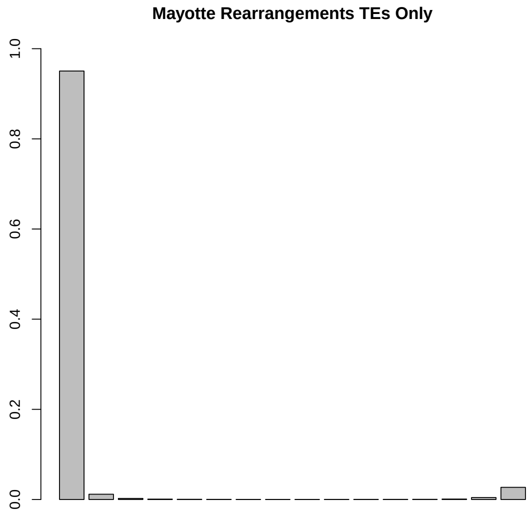


A

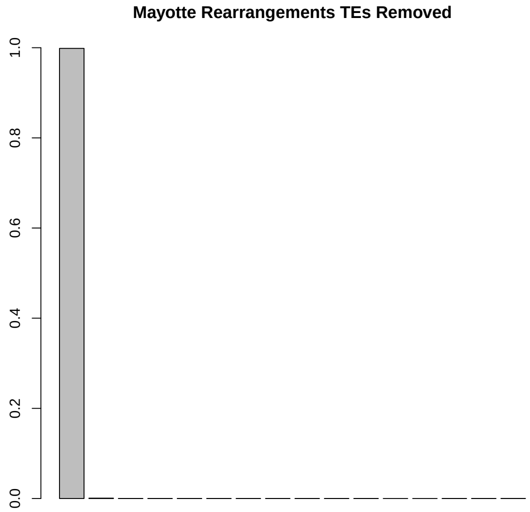


B

Figure S11: Derived allele frequency distributions are shown for rearrangements associated with transposable elements (TEs) and those without TE associations. (A) TE-associated rearrangements exhibit a skew toward higher allele frequencies, consistent with increased retention or possible selection acting on TE-mediated variants. (B) Non-TE-associated rearrangements are more uniformly distributed across frequency bins, suggesting different underlying mutational or selective dynamics. Overall, 57.9% of rearrangements in Mayotte were associated with TEs, indicating that TE activity plays a major role in shaping structural variation in this population.
